# Standardizing a Protocol for Studying pH-dependent Transient Conformations using Computer Simulations

**DOI:** 10.1101/2020.12.21.423456

**Authors:** Ruhar Singh, Andrew M Lynn

## Abstract

Understanding pH-dependent protein stability is important in biological - transport, storage, and delivery, i*n vivo* conditions such as aggregation potential in neurodegenerative disease, and in studying the folding/unfolding of proteins. Using computer simulations, we can replace complex experimental determination and provide an atomistic-level interpretation of the cause and effect of pH on protein stability. Here, we standardize a method that provides a framework through which we examined pH-dependent transient conformations during unfolding simulations of proteins. Constant pH simulations utilized in the prediction of pKa values of charged groups of the peptide. The calculated pKa values employed to fix the appropriate protonation state of the amino acid to simulate the effect of pH on the system. Trajectories from multiple high-temperature MD simulations of the protein sample the conformational space during unfolding for a given pH state. The ensemble of conformations is analyzed from its free energy landscape to identify transient and stable conformations both at a given pH and between different pH. As a test system RN80, a protein fragment analog of the C-peptide from bovine pancreatic ribonuclease-A used to measure the accuracy of the predictions from simulations. Experimental measures of the helix content determined as a function of pH display a bell-shaped curve, i.e. RN80 alpha-helix formation is maximum at pH5 with a subsequent loss in helicity at higher and lower pH. The main forces stabilizing the alpha-helix are a salt-bridge formed between Glu-2 and Arg-10 and cation-pi-interaction between Tyr-8 and His-12. Our protocol includes constant pH calculations, optimal high-temperature simulations, and Free Energy landscape analysis exhibited the agreement with the experimental observations.

## 2. INTRODUCTION

Protein folding is a process of molecular self-assembly in which the protein collapses to form a compact well defined three-dimensional structure in the native state[(Dill et al. 2008) ref]. Protein folding studies propose that helix formation occurs in the early stage of folding, which after that coalesces to form tertiary structure[(Shea and Brooks 2001; Bierzynski, Kim, and Baldwin 1982; Brown and Klee 1971)]. The C-peptide from bovine pancreatic ribonuclease is 13 fragments long peptide that shows alpha-helical conformation in an aqueous solution. Recently [(Jesus et al. 2018; Sugita and Okamoto 2005)] have demonstrated that specific interactions such as salt bridge (Glu-2 and Arg-10) and cation-pi interaction (Tyr-8 and His-12) play a crucial role in the stability and folding dynamics of helical peptide[(Shoemaker et al. 1987)]. NMR and CD spectroscopy revealed that C-peptide has a three-state pH-induced conformation and the high peak in the curve is displayed by RN80 alpha-helix formation with maximum helical content at 3°C and pH 5 which decreases on moving towards acidic or alkaline pH (Figur1(a))(Jesus et al. 2018). At decreasing pH, the protonated form of His-12 favours the folded conformation through the cation-pi interaction although the Glu-2 protonation induces partial unfolding due to the dissociation of the salt-bridge (Fairman et al. 1990).

**Figure1:**
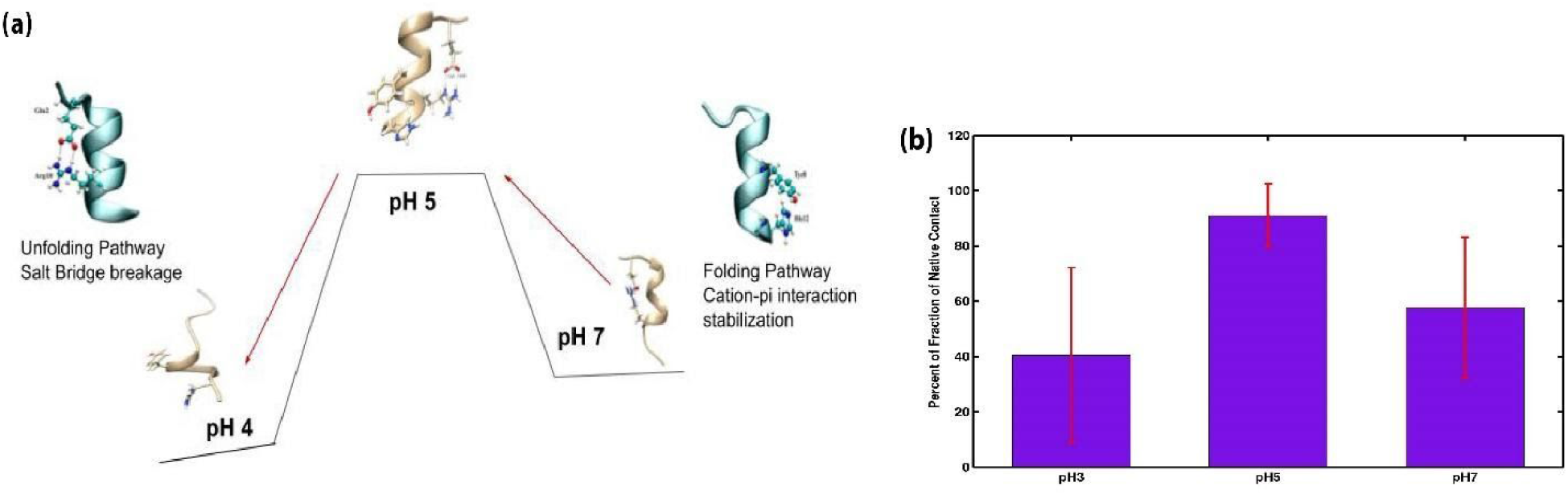
(a) Experimental verification of C-peptide stability at different pHs. (b)The fraction of native contact averaged over three replicates for pH3, pH5, and pH7 respectively, at 276K temperature.

**Figure2:**
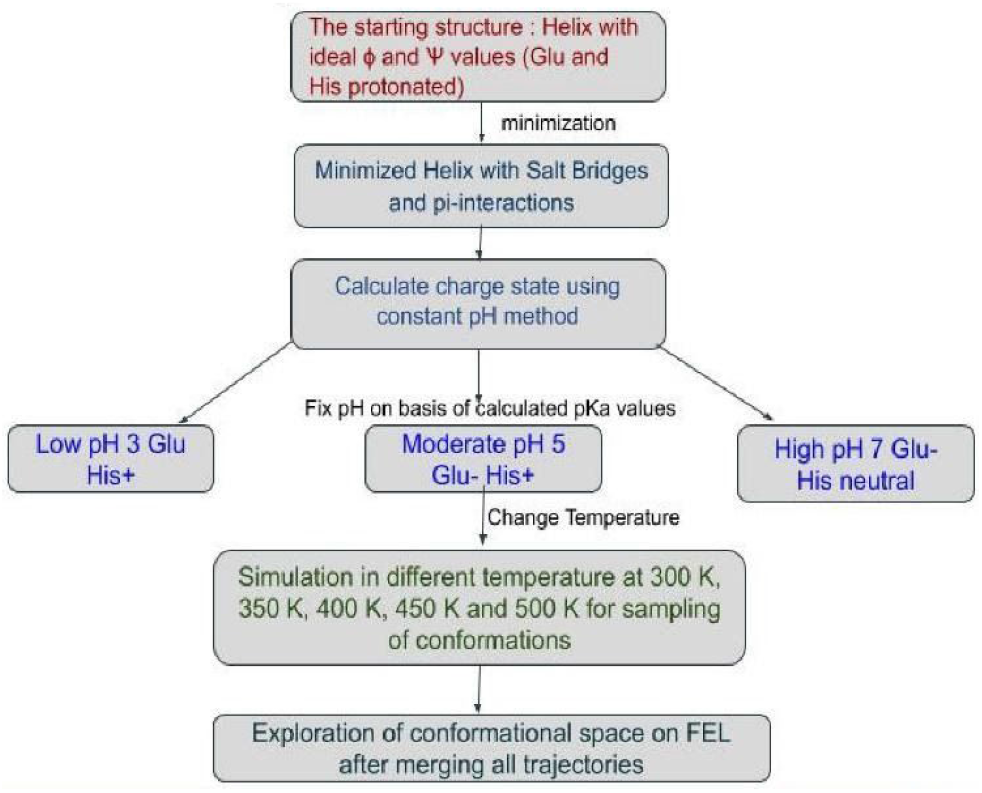
Methodology for the generation of structure and setting up a simulation.

Here, we established a protocol that provides a general approach and standard setup for the MD simulation to study the pH-dependent behaviour of proteins. The currently established protocol is validated with C-peptide as a model system and perform simulations at various conditions of pH and temperature. It would capture the transitional and significant conformation of the peptide on a free energy landscape. The structural change in C-peptide (RN80) is characterized through conformation, energy, and dynamical parameters at different pH. It would visualize the impact of pH on protein conformation and charged residues on salt-bridge and cation-pi interactions. By overlaying conformational sampled space derived from trajectories at individual pH on a standard free energy surface, it is possible to predict pH-dependent transitions using Markov state models. pH-dependent states involve protonation or deprotonation, which is not possible to simulate using classical molecular dynamics. The alternative methods of the currently proposed protocol are REMD and Thermodynamic integration which are used for exploration activities of conformation space, transition and free energy calculation.

## 3. DEVELOPMENT OF PROTOCOL

### For interpretation thermodynamic integration from free energy plots using small peptide

#### Initial structure

C-peptide ribonuclease (RN80 ribonucleaseA) is a 13 residue long helical peptide with sequence AETAAAKYLRAHA contained blocked ends with potential salt bridge and cation-pi interaction (Glu2-Arg10 and Tyr8-His12 with i to i+4).

##### i. Calculation of pKa values using Constant pH MD method and Generation of pH specific structures

The constant pH simulation method enables molecular dynamics simulation to spontaneously change the protonation states of the bio-molecular system during the simulation (Dobrev et al. 2017). The pKa values of ionizable groups calculated from the distribution of the protonation states(W. Chen et al. 2014) shown in figure S1. The calculated pKa values for charged residues are the following: LYS 9.92, Tyr 10.35, His 6.41, and Glu 4.89, respectively. The initial model of the peptide built as an ideal α-helix with extended side chains. The charge state of peptide was fixed on the basis of pKa values. The ionization states of the amino acids represent the peptide in three pH regions: low pH (e.g., both the Glu and His residues were protonated correspond to <pH 4), moderate pH (Glu negatively charged and His protonated, pH 5), and high pH (Glu negatively charged and His neutral with proton on N corresponding to < pH 7 (≈ pH7))

##### ii. Sampling of conformations at a different temperature and Molecular dynamics simulations of C-peptide

There are numerous methods established for the sampling of the protein conformations and explore the conformational space such as replica-exchange accelerated MD and high-temperature simulations. Here, the most beneficial point of using temperature is crossing the barrier in the free energy landscape, so the unfolding rate will increase. The high temperature and non-covalent interactions such as cation-pi and salt-bridge were used for space sampling and constraints respectively for characterization of conformational transition of C-peptide. We performed simulations at three pH (3, 5 &7) states using various temperatures (276K - 500K) to explore the conformational space of C-peptide in different conditions.

##### iii. Overlay of conformational states on common FEL

Markov state model (MSM) has been used in the analysis of protein dynamics as it provides a framework to study the conformational transitions occurred during the simulation[(Husic and Pande 2018)]. The transition state and folding path of multiple simulations can be predicted. The order of structural element formation/breakage can be used to quantify the heterogeneous folding events in transition paths of different trajectories.

### 3.1 We restrained the system for the stabilization of the Cation-pi interaction

AMBER and available force field were not able to reproduce the cation-pi interactions, to the levels expected from experiment. We put cation-pi interaction by fixing an anion (Fl) at the center of the aromatic ring[(Dobrev et al. 2017; Hanif M. Khan et al. 2016)] which modifies the LJ (Lennard Jones) parameters or vdw interactions (Figure3). This helps mimicking the cation-pi interaction on C-peptide between Tyr8 and His12, by providing an attractive force on the Tyr ring for the partial charge on Histidine (Orabi and Lamoureux 2018).

**Figure3:**
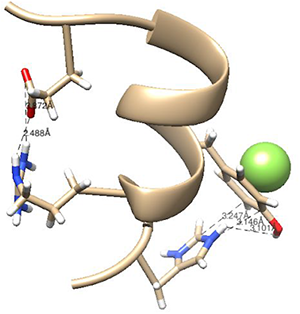
The native structure of C-peptide with cation-pi (Tyr8-His12) and Salt-bridge(Glu2-Arg10) interactions which is restraint with Fl-ion for capturing the cation-pi interactions.

### 3.2 Verification of cation-pi and salt-bridge interactions in C-peptide structure

We simulated the C-peptide at different conditions of 276 and 300K temperatures, vacuum under three pHs 3, 5, and 7 respectively (Figure S2 and S3). Native contacts show that the C-peptide is significantly stable at pH 5 and the distance between salt-bridge forming residues is maintained at moderate and high pH, while large excursions are observed at low pH. Cation-pi interaction proved to be a poorly stabilizing factor (FigureS2(b)). After performing restrained molecular dynamics simulation, measured the distance between cation-pi forming residues. The deviation in distance of cation-pi residue is less in a restrained condition which shows that it is well maintained. The modified parameter set results in cation-pi interaction matching with experimental data. FigureS3(b) demonstrates the cation-pi (His-Tyr) and salt-bridge (Glu-Arg) distance.

## 4. MATERIALS AND METHODS

First, generate a 13-residue fragment of C-peptide ribonuclease (RN80 ribonucleaseA) with blocked terminal residues which contains the ideal value of ϕ and ψ of an α-helix, after which minimization performed so that the stable cation-pi and salt-bridge interactions can be formed. Then, the pKa values of titratable residues are estimated using the constant pH method. Furthermore, the conventional molecular dynamics simulation has been performed. The charge state of the residue has been fixed on the basis of pKa values to the corresponding pH in simulation. The peptide was explicitly described by employing an all-atom model. The solvation accounted for the solvent model TIP3P water and set the box edge side as 8 Ang in which the minimum distance is 1 Ang. The entire system was minimized with 500 steps of steepest descent followed by 500 degrees of conjugate gradient before the production run. In the heating phase, the solute was harmonically restrained with 10 KJ.mol^−1^. nm^−2^ performed for 100 ps which followed by the 500 ps equilibration phase without restraints. All MD simulations performed with varying conditions of pH and temperature. Molecular dynamics simulation tool, Amber16 was used to carry out simulations at different temperatures; 276K, 300K, 350K, 400K, 450K, and 500K under various pHs. We employed MSMBuilder package for conducting most of the investigation in IPython scientific environment with the use of the matplotlib and scikit-learn libraries, images were generated by chimera and secondary structure analysis performed by using DSSP program.

## 5. RESULTS

We used our standardized protocol to generate trajectories for exploration of conformational space and deduction of the folding pathway at different pH and temperature, from this observed various events of folding and unfolding. The multiple trajectories are generated of 1 ns simulation from low to high temperature at various conditions of 10 replicates started from three pH conformations. Here we obtained all the possible conformations of C-peptide from native to extended conformations after merging all the simulation data. Altogether, from this way, collected the dataset which consisted of 180 trajectories (6 temperatures X 3 pHs X 10 replicates) in various conditions and then analyzed by MSM method to calculate the free energy landscape and quantify the accessible states of C-peptide in low energy basins.

### 5.1 A sampling of conformations at different conditions and DSSP allows the observation of the evolution of change in the secondary structure during the unfolding

In the current approach, we are investigating the conformational transition of the C-peptide at various conditions including high temperature and pH. We run numerous MD simulations and generated 1ns simulation trajectories with 180ns aggregated simulation time for enrichment of the conformational space followed by various temperatures such as 276, 300, 350, 400 450, and 500K over 10 replicates in all pH3, 5, and 7. We sampled the conformations and gain detailed information regarding secondary structure change for this calculated DSSP plot, as shown in figure4. All the secondary structures were stable at low temperature but as moved towards the high temperature (<350K, 400K, 450K, and 500K) the secondary structure start melting. Figure4 is showing that initiation of simulation during equilibration and low temperature from which observed the data-rich regime of secondary structure. However, the helix broke up as a result of the secondary structure declined indicates the importance of interaction in the stabilization of C-peptide structures. The DSSP plot visualizing the secondary structure change in multiple conditions although linked with occupied conformations in the basin on the free energy landscape.

**Figure4:**
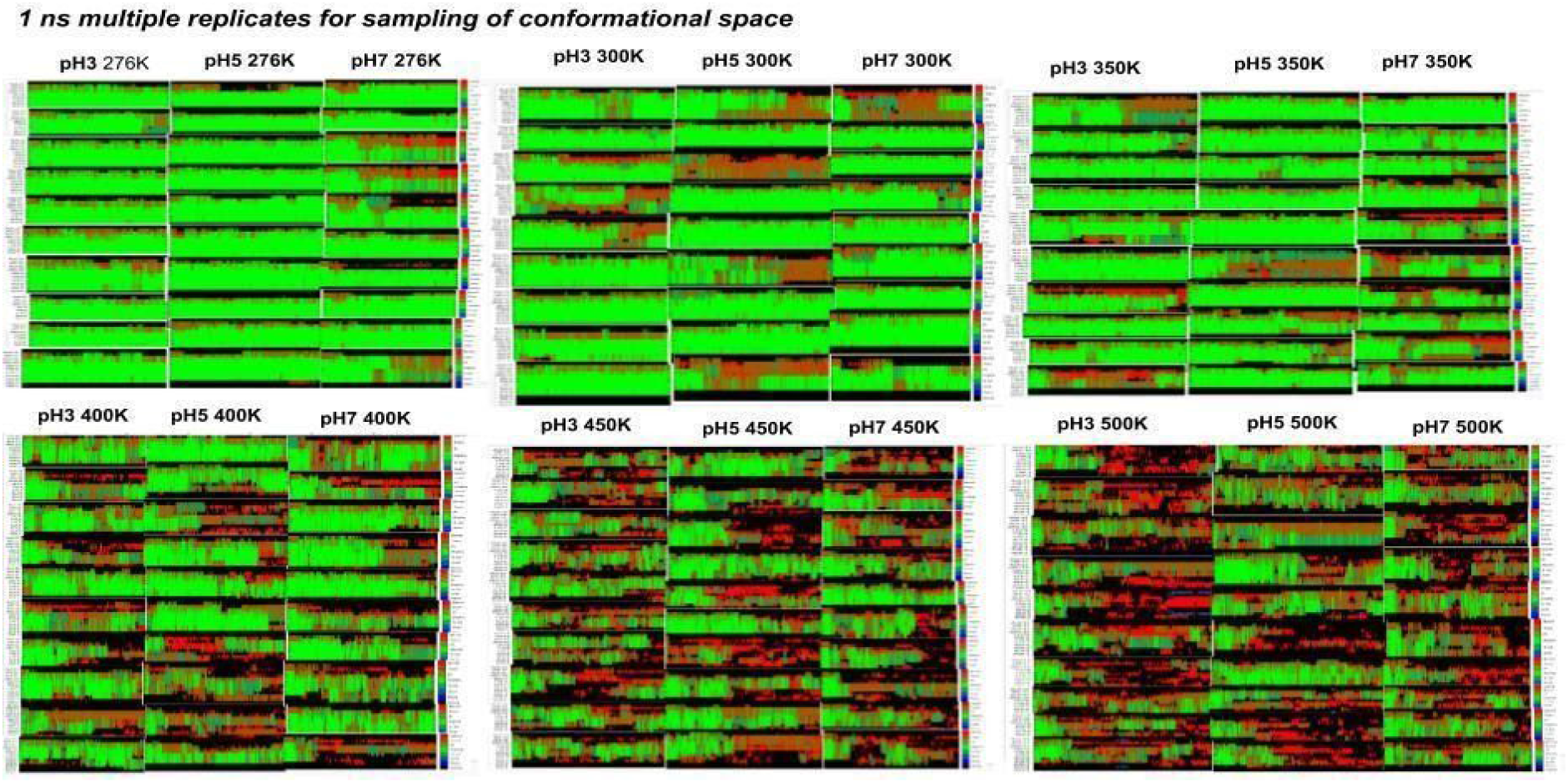
The change of secondary structure concerning time evolution of C-peptide calculated from unfolding simulation using sampling at pH3, pH5 and pH7 and various temperatures 276K, 300K, 350K, 400K, 450K, and 500K respectively, the helix is marked in green color and bend and turn in red color.

### 5.2 Quantification of the conformations on grid points calculated from the free energy landscape

MSM-Builder relates the structural behavior of C-peptide in various conditions through free energy surfaces and the underlying process is the Markovian. We used enhanced sampling methods such as high temperature for conformational sampling in reduced time scales which distribute configurations from low energy states to the high energy states and assign populations at different energy basins. It recovered the canonical distribution and Boltzmann ensemble of all populations on the free energy. The sampled trajectories are projected onto the two dominant slow processes of tICs from the tICA model on a free energy landscape (Figure5(a)); the dominant tIC1 values are correlated with positive or near zero value of tIC2 to low energy configuration on FEL corresponding to the various pH. The range lies on FEL for tIC1 is 0.95 to −4.85 and for tIC2 is −5.10 to 4.20 and the centrally located region ranges from 0.95 to 0.37 at tIC1 and - 0.45 to 0.48 at tIC2 on FEL. We built a grid of 10×10 along the free energy surface, applied the histogram2d and calculated the population at each grid corresponding to the basin. Then 2D histogram helps in count number quantification or estimation of the number of conformations in a bin as shown in figure5(b). However, the calculated population is the measurement of conformational density occupied into various energy basins. At central basin, we have seen that maximum conformations are confined to low energy states with around 7500 conformations at 276K for all pH, after that, about 1000 conformations into second column basin mostly at 300K temperature, and here it started to involve with an inlet at high energy states. Further, basins occupied number of conformation is 500, 200, 100, 50 and 25 as moving towards at high temperature 350K, 400K, 450K, and 500K temperature and also split into three pH region on FEL. pH7 are confined into the upper region, pH5 into the middle region and pH3 onto the bottom region of FEL respectively. As moved from left to right mostly folded conformation at low temperature, then in the middle region various intermediate present at different energy levels and lastly after the 5th column primarily unfolded conformation were seen. The centrally located basin contains maximum conformation which belongs to native conformation of pH3, 5, and 7. The overall observation found that as moving towards a high temperature, the conformation populations are sequentially decreased and the conformations occupied high energy state. We obtained information on the distribution of conformation along with energy levels.

**Figure5:**
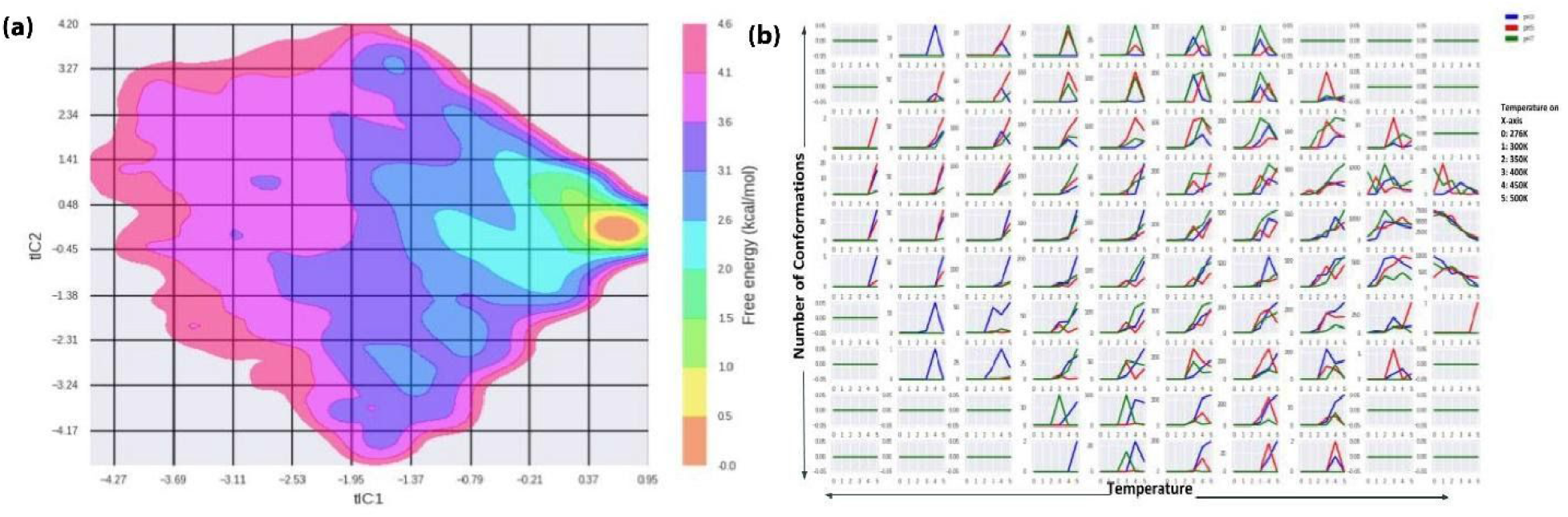
(a) Projection of dihedral angle of C-peptide conformations in different pH states onto free energy landscape calculated from unfolding simulation from sampling at pH3, pH5 and pH7 and various temperatures 276K, 300K, 350K, 400K, 450K, and 500K respectively. The calculated free energy map is shown as a contour plot in a grid. (b) The quantified population of conformations of C-peptide in the energy basin calculated from histogram binning of the free energy landscape of the three pH states which are separated by temperature. The pH5 populations are marked red, pH3 populations in blue and pH7 populations in green color respectively.

**Figure6:**
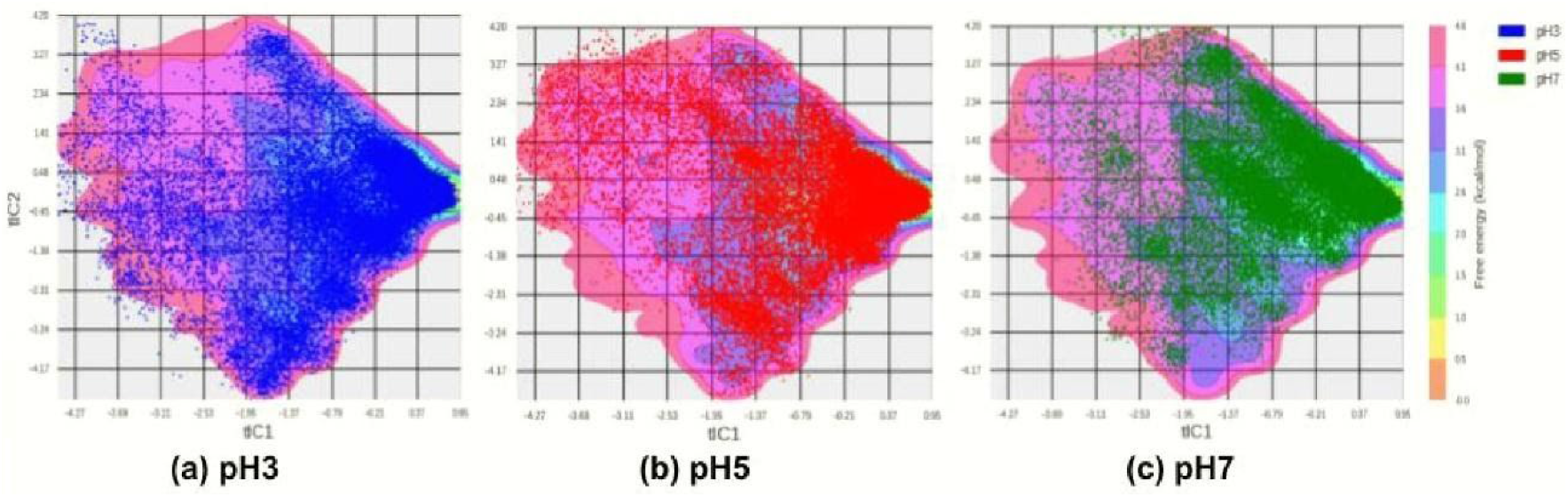
The population density corresponding to each pH is calculated and overlaid onto the free energy landscape which represents the trajectories movement onto the phase space the (a)pH3, (b)pH5 and (c)pH7 conformation are marked red, pH3 conformation in blue and pH7 conformation in green color.

### 5.3 A density plot segregating contribution to the free energy landscape by pH and temperature can show the path of the unfolding

Here, the trajectories are overlaying onto the free energy landscape for the interpretation of the molecular movement or direction of trajectories corresponding to different pH. It also provides the visual intuition of the state conformation, which is going from one part of the phase space into another. We observed that the maximum overlap of pH3, pH5, and pH7 trajectory into low energy basin at 276K temperature after that as move toward high temperature generated trajectory the conformations are also split into pH3 state, pH5state and pH7 state occupied region on FEL. The primary form of conformation which has both interactions salt-bridge and cation-pi remains stable during the early phase of protein unfolding and occupied overlapped low energy basin which correspond to conformations of pH3, pH5 and pH7 and retained some secondary structure for those states. During the later stage, the secondary structure started to decline and occupied those three stages which describe the three conformational states corresponding to pH3, pH5, and pH7.

### 5.4 Markov State Models can be used to recreate the free energy landscape after Merging the individual unfolding trajectories

For the detailed analysis of the molecular mechanism of the folding and stability of C-peptide applying the free energy landscape as shown in figure7. A set of simulations performed to sample the conformational space, and it will provide the average structure properties. Here the proposed protocol overcomes the limitations of the established simulation method by enumerating all possible unfolded conformations and predicts the accurate properties. The structure changed with several pH imposed conditions which is not changing on a single trajectory simulation. The low energy basin contains so many conformations which are reappeared many times in the MSM trajectory, which means that some conformation which includes both interactions are high in number. These conformations within the trajectory are visited several times, and going in and out from the microstates. The highly populated conformations correspond to local minima in the free energy landscape, which is the most stable structure. The conformational transition from α-helix (F) to coil form (U), giving an idea that “helix unwinding” in which these macrostates are representing the intermediate states during transition. By following the protocol, we predicted three conformations which resemble three pH5, pH3, and pH7 conformations after sampling of large conformational space. Although the results showing in figure7, the termini are stabilized due to the cation-pi interactions and contains distance 3.04 Å in microstates 9 at pH3 and observed the salt-bridge interactions with length 4.02 Å in microstate 3 which causes peptide to remain intact and stable during the simulations at pH7. The overall picture reveals that alpha-helix is maximally stabilized at pH5 which contains both interactions cation-pi and salt-bridge with 3 and 4 Å distance respectively. The set of conformational states and the transition probabilities between pairs of states showing possible pH state conformational change of C-peptide on the free energy landscape. The transition probability matrix between states to identify the possible pathways among states which shows that pH3 and pH7 representative conformations cross the high energy barrier and move towards the low energy basin. States 0 and State 1 corresponding to pH3 and pH7 conformations are in high energy states 2.6 and 2.0 kcal/mol with transition probability of 0.156 and 0.154 goes towards microstate2, figure7(ii) depicted that they contain the high probability to move towards pH5 states corresponding to low energy basin (0-0.5 kcal/mol). The transition occurs pH5 conformations to pH3 and pH7 conformations on the free energy landscape in starting and further its move towards the unfolded state. From this also predicted possible transition of conformation from pH5 state to pH7 state and pH5 state to pH3 state, the whole procedure termed as Thermodynamic Integration. Here quantified the conformation on FEL and found that pH5 state contains maximal conformation after that pH3 state and pH7 state confined its conformation and also found out the possible transition between conformation. It indicates that the states, containing similar secondary structures, are interconverted rapidly on the free energy landscape This fundamental analysis gives insight into how C-peptide gradually loses the helical character and occupies the different energy basin configurations also shows the possible transition among pH states of the peptide.

**Figure7:**
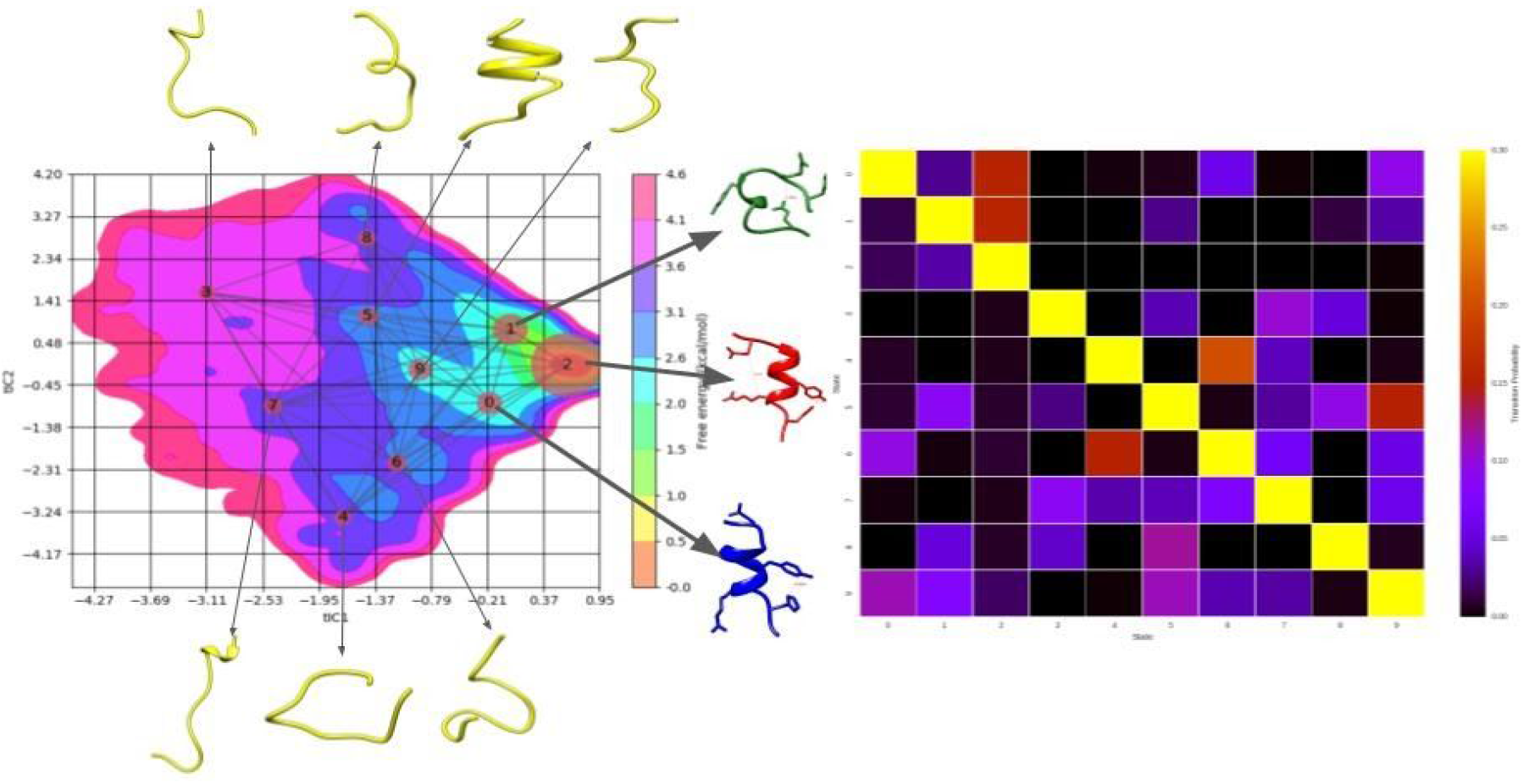
MSM free energy map and flux pathway draw from unfolding simulation using sampling at pH3, pH5, and pH7 and various temperatures 276K, 300K, 350K, 400K, 450K, and 500K respectively. In the 10-state microstate model, the circle sizes representing the relative populations of each microstate which contribute to the transition from a fully folded state (state 0) to a fully extended state. The pH5 conformations are marked red, pH3 conformation in blue and pH7 conformation in green color. (ii) heatmap representation of transition probability matrix to visualize the possible transition between states.

## 6. DISCUSSION

This study characterizes the kinetics and folding-unfolding of C-peptide α-helix in which two protonated amino acids Glu-2 and His-12 play a significant role in the stabilization of helix. At pH5 besides canonical hydrogen bonds, the salt bridge between Glu-2 and Arg-10 and cation−π interaction involving His-12 and Tyr-8 contribute to the stabilization of α-helical conformation. Here, implemented a protocol using different temperatures and pHs to validate the computationally intensive sampling of conformational space effectively. The data simulated at various conditions, merged all the trajectories, clustered the conformations and identified occupied states on the free energy landscape. After conformational sampling at various conditions the long time period simulation also performed with three replicates in three pHs for verification of experimental measurements shown in figure1. We observed that the cation-pi distance is not maintained - there is no explicit parameter for this interaction in the force field, so we fixed the anion (Fl) towards the center of the aromatic ring perpendicular to the Tyr aromatic ring for modifying the explicit electrostatic interactions. Therefore, the anion was restrained between the Tyr8 and His12, to capture the cation-pi interaction. Fl is highly negative charge atoms, it contains high electrostatic contributions in stabilization and also influences other charged residues. But here not considering the energetic involvement only quantified the structural parameters based on dihedral angles. Thus, not examined as a significant impact on results. Although, standardization of the partial charges on anion atoms required to maintain the cation-pi interactions without influencing the salt-bridge interactions is a possible solution.

The conformational landscape links the various folded and unfolded states which are connected by structurally diverse intermediates. Further, quantified the states corresponding to pH3, pH5, and pH7 into the numbers. Through the comparison of count values in different pH also speculated the propensity of the conformations which underwent folding/unfolding and clustered the conformation into various energy basins, which reveals the unfolding pathways of various pH generated trajectories with different probability of the macrostate. Here, the proposed protocol includes enhanced conformational sampling and recovering of the canonical ensemble. We merged the multiple trajectories generated at multiple temperatures and computed the free energy landscape. The unweighted FEL has been calculated as the coverage of conformational space and well depth is not the primary aim of study. The FEL recovered from the current implementation is without population reweighting. The weighting of populations can be performed by using a weight based on temperature to the conformer populations calculated using the Boltzmann distribution for retrieving the corrected number of conformations in each energy basin. Further, here, find the population of pH states in the energy basin, kinetics of their interconversion, and protonation state on the corresponding pH, which establish possible potential of MD and MSM technique which effectively predict the protonation state change. pH3 and pH7 macrostates goes towards pH5 macrostates without breakage of bond. pH5 conformations were thermodynamically stabilized and have been in a deep energy basin constructed from MSM. It provides a full description of the protonated and deprotonated state conformational movement and possible transition.

## 7. CONCLUSION

Here, carried out an MD simulation of 13-amino acid long C-peptide α-helix at various temperatures and pHs. After merging all trajectories, identified conformations belonging to different pHs during the transition from unfolding to folding state. It characterizes the stability and transition of conformations relied on free energy. From this, predicted the pH-dependent transitions (protonation and deprotonation) on free energy surface using MSM approach which is not feasible with the conventional molecular dynamics simulation. By following the protocol, speculated three conformations which resembled three pH 5, pH3, and pH 7 after sampling of large conformational space. The information about the distribution of conformations along the energy levels are obtained from the count numbers and also plotted along the free energy landscape. The interconversion of multiple states which are occupied at different pH in different energy levels are calculated on the basis of their structural similarity on the FEL. This whole procedure we named thermodynamic integration through this, calculate the consecutive conformations at various pH states and also properties at the various energy levels. Overall method clarifies the folding mechanism and occupied states on the FEL.

## 8. Scripts and Dataset Availability

The scripts and datasets are publicly available on GitHub.

**GitHub URL: https://github.com/jnu63/cpeptide-msmcode**

## 9. Acknowledgement

RS gratefully acknowledged to funding agency Department of Science and Technology (DST/INSPIRE Fellowship/2015/IF150960), Govt. of India and JNU for financial support and fellowships. The authors acknowledge the Computational Facility at SCIS, JNU.

## Notes

### Competing Interest Statement

The authors have declared no competing interest.

