## Supplementary Figures for "Standardizing a Protocol for Studying pH-dependent Transient Conformations using Computer Simulations"

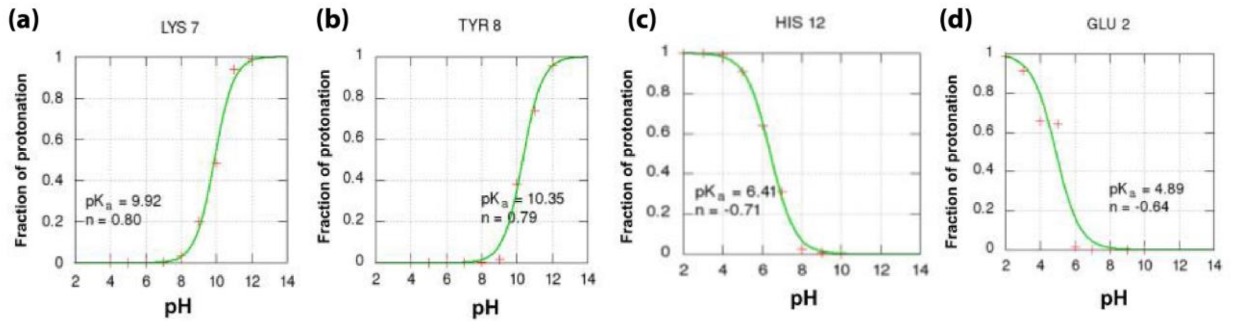


**FigureS1: Titration curves of acidic and basic residues (a) Lys7 (b)Tyr8 (c) His12 (d)Glu2. The residues are analyzed separately. The curves were fitting with the procedure. (Constant pH simulation results)**

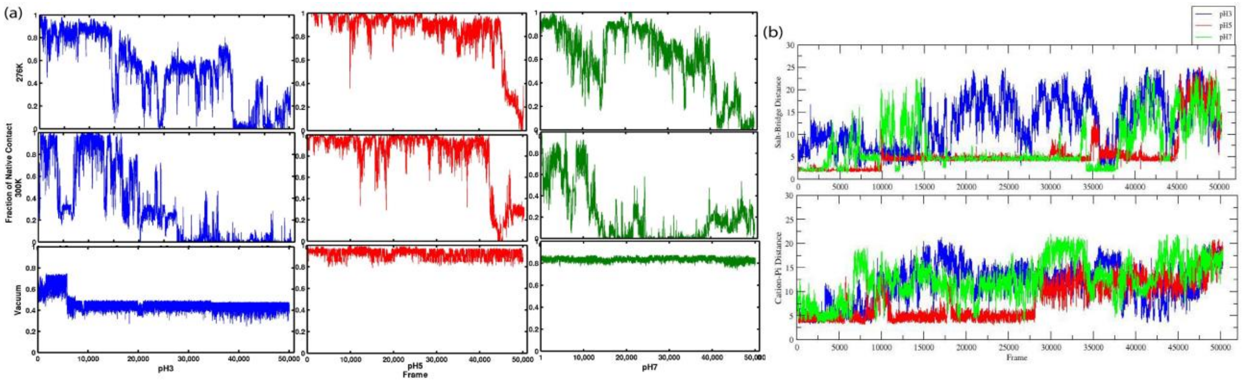


**FigureS2: (a)Fraction of Native Contact with time evolution shown for all the simulated systems at 276K, 300K and vacuum for pH3, pH5, and pH7 respectively corresponding to without restrain condition. (b)The salt-bridge and cation-pi distance as a function of simulation time. The different pH 3, 5, and 7 represented by colour code blue, red, and green respectively.**

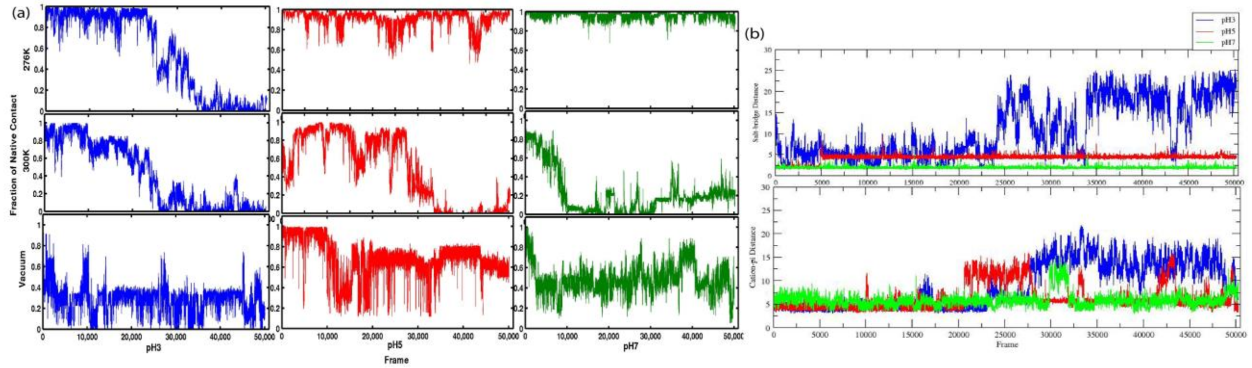


**FigureS3: (a)Fraction of Native Contact with time evolution shown for all the simulated systems at 276K, 300K and vacuum for pH3, pH5, and pH7 respectively corresponding to the restraint condition. (b)The salt-bridge and cation-pi distance as a function of simulation time. The different pH 3, 5, and 7 are represented by colour code blue, red, and green respectively with the restraint condition.**
